# Analytical code sharing practices in biomedical research

**DOI:** 10.1101/2023.07.31.551384

**Authors:** Nitesh Kumar Sharma, Ram Ayyala, Dhrithi Deshpande, Yesha M Patel, Viorel Munteanu, Dumitru Ciorba, Andrada Fiscutean, Mohammad Vahed, Aditya Sarkar, Ruiwei Guo, Andrew Moore, Nicholas Darci-Maher, Nicole A Nogoy, Malak S. Abedalthagafi, Serghei Mangul

**Author notes:** These authors jointly supervised this work (Nicole A Nogoy, Malak S. Abedalthagafi, Serghei Mangul).

## Abstract

Data-driven computational analysis is becoming increasingly important in biomedical research, as the amount of data being generated continues to grow. However, the lack of practices of sharing research outputs, such as data, source code and methods, affects transparency and reproducibility of studies, which are critical to the advancement of science. Many published studies are not reproducible due to insufficient documentation, code, and data being shared. We conducted a comprehensive analysis of 453 manuscripts published between 2016-2021 and found that 50.1% of them fail to share the analytical code. Even among those that did disclose their code, a vast majority failed to offer additional research outputs, such as data. Furthermore, only one in ten papers organized their code in a structured and reproducible manner. We discovered a significant association between the presence of code availability statements and increased code availability (p=2.71×10^−9^). Additionally, a greater proportion of studies conducting secondary analyses were inclined to share their code compared to those conducting primary analyses (p=1.15*10^−07^). In light of our findings, we propose raising awareness of code sharing practices and taking immediate steps to enhance code availability to improve reproducibility in biomedical research. By increasing transparency and reproducibility, we can promote scientific rigor, encourage collaboration, and accelerate scientific discoveries. We must prioritize open science practices, including sharing code, data, and other research products, to ensure that biomedical research can be replicated and built upon by others in the scientific community.

## Introduction

Modern biomedical research relies heavily on quantitative analysis, necessitating mechanisms that ensure transparency, rigor, and reproducibility. Since 2011, an alarming number of retracted papers have been quantified in a “retraction index” indicating a strong correlation with journal impact factor^1^. This “retraction index” highlighted weaknesses in the publishing system^2^, further fueling a lack of trust in science. Twelve years after the “retraction index”, the number of papers being retracted has increased, from 45 per month in 2010 to 300 retractions per month in 2022, reported by Retraction Watch^3^. Researchers have a responsibility to help mitigate such problems, by carrying out best practices and doing better science - ensuring transparency and reproducibility of their work and increasing trust.

Transparent and reproducible research requires the sharing of well-documented analytical code and data. However, analytical code sharing is not enforced as strictly as data sharing^4,5^, and the effectiveness of existing initiatives is uncertain. Well-structured data and code not only enhance reproducibility, but also provide numerous benefits to researchers--they facilitate swift installation and rerun of analyses, enable visualization, and make analyses more robust against code errors and other mistakes^6,7^. By contrast, the limited availability of code and associated research products poses challenges to reproducibility across scientific disciplines^8,9,10,11^. While funding agencies and peer-reviewed journals have recommended best practices, guidelines, and checklists for researchers, including for sharing of analytical code^8,12^, these recommendations often have little impact on the actual availability of research products accompanying research studies^8^. This situation is present despite the rise of data-driven biomedical research, and the growing recognition within the broader scientific community about the significance of sharing code used for analysis^13^.

Our study aimed to assess the code sharing patterns and practices in biomedical research by analyzing 453 manuscripts published in eight scientific journals between 2016 and 2021. Our analysis suggests that the majority of the surveyed studies did not share analytical code used to generate results and figures. Furthermore, among the subset of studies that did share their code, a substantial majority did not share other research outputs. Furthermore, only a small number of studies organized their code in a structured and reproducible manner. We discovered a significant association between the presence of code availability statements and increased code availability (p=2.71×10^−9^). Out of the 453 manuscripts analyzed, 43% contained code availability statements, and 85% of those manuscripts shared code. In contrast, only 10% of the manuscripts that did not declare code availability actually shared their code. Our study also revealed an increase in code sharing practices from 41% in 2016 to 65% in 2021, while raw data sharing practices increased from 24% in 2016 to 38% in 2021. In light of our findings, we propose raising awareness of code sharing practices and taking immediate steps to enhance code availability to improve reproducibility in biomedical research. By increasing transparency and reproducibility, we can promote scientific rigor, encourage collaboration, and accelerate scientific discoveries. We must prioritize open science practices, including sharing code, data, and other research products, to ensure that biomedical research can be replicated and built upon by others in the scientific community.

### The majority of biomedical studies has limited code availability

In 2015, a Repeatability in Computer Science study was carried out to see the extent to which Computer science researchers share their source code^14^. Eight years later, we conducted an extensive study to assess the availability of code and data in biomedical research, analyzing a random sample of 480 articles across eight biomedical journals published from 2016 to 2021 (Table 1). Ten manuscripts were selected from each journal for each year of the study period. Out of 480 manuscripts, we considered 453 manuscripts for further analysis. The remaining 27 manuscripts were research letters, editorials, or research reports which did not report any data or code and were discarded from the analysis. In our analysis, we recorded code availability, data availability in the manuscripts, and the presence of data and code availability statements and code organization. Our results indicate that nearly half (49.9%) of the studies examined failed to share the analytical code used to generate the figures and results used by the researchers (Figure 1a). Of the code that was shared, the majority (76.92%) was found on GitHub, while 13.66% was hosted on other platforms such as Bitbucket, Sourceforge, or individual websites, 5.98% on Zenodo, 2.56% was included as supplementary material, and 0.85% on GitLab (Table 1). Of the 453 studies, 209 (46.14%) used a code availability statement, while 179 (85.65%) of these 209 actually shared their code (Figure 1b). However, even if the paper did share the analytical code, there was still the question of whether the link for the code itself functioned or if it had an archival stable link. The term “archival stable link” refers to a permanent or persistent link that remains functional and accessible over time, ensuring that it continues to point to the same resource or information even if the website structure or content undergoes changes. To assess the archival stability of code sharing repositories, we examined the links of the manuscripts that did share the code. Surprisingly, we found that 1% of the links were unstable, indicating potential issues with long-term accessibility.

**Table 1:** Code and data sharing policies across 8 biomedical journals

|  | Journal | Code Policy | Data Policy |
| --- | --- | --- | --- |
| **0** | Bioinformatics | Mandatory | Mandatory |
| **1** | BMC Bioinformatics | Encouraged | Encouraged |
| **2** | Genome Biology | Mandatory | Mandatory |
| **3** | Genome Medicine | Encouraged | Encouraged |
| **4** | Nature Biotechnology | Mandatory | Mandatory |
| **5** | Nature Genetics | Mandatory | Mandatory |
| **6** | Nature Methods | Mandatory | Mandatory |
| **7** | Nucleic Acids Research | Encouraged | Mandatory |

**Figure 1.**
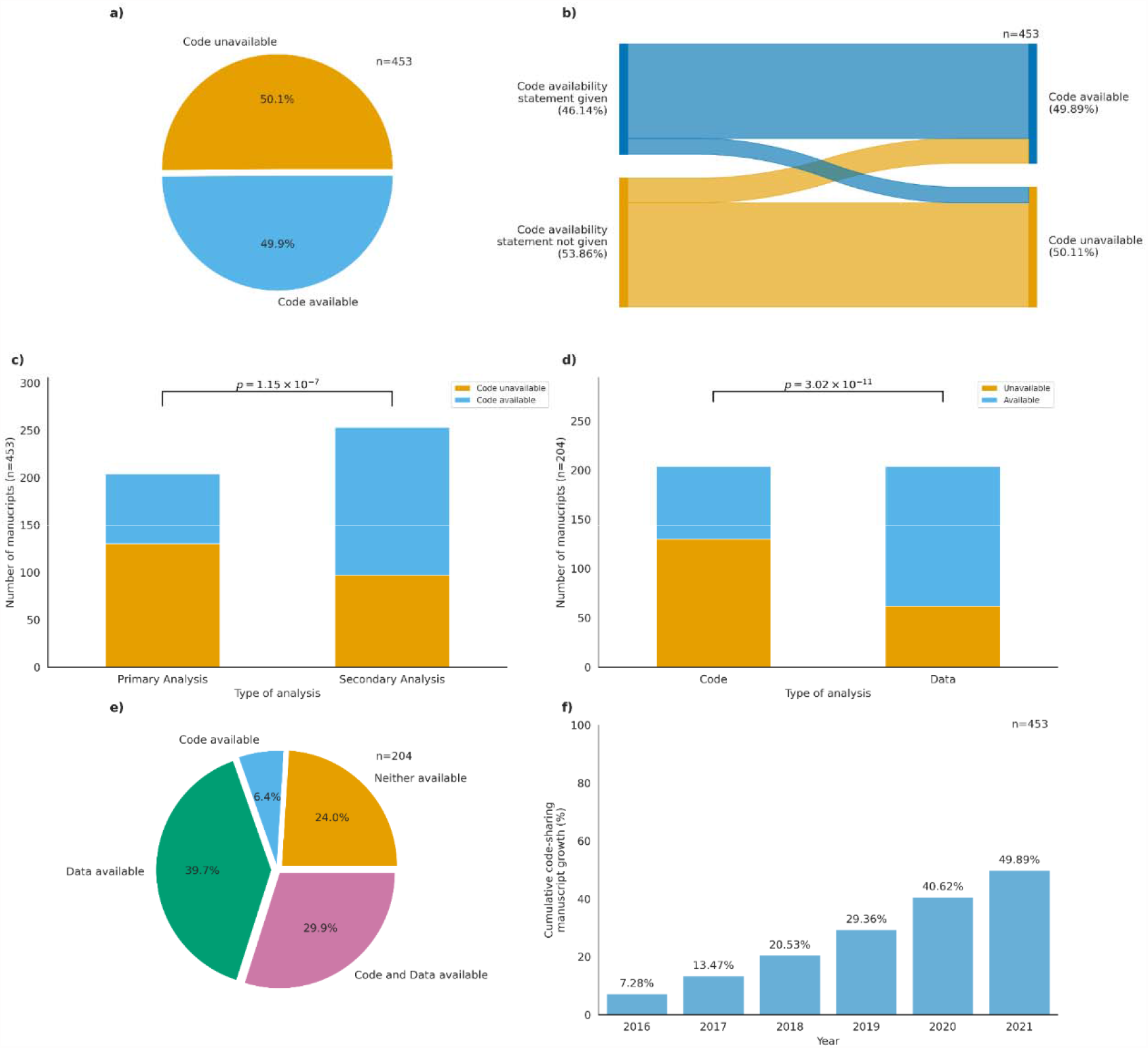
**(a)** Availability of code across the 453 biomedical articles. **(b)** Code availability status across 453 manuscripts on the basis of code availability statement. **(c)** Number of studies sharing code across primary and secondary analysis studies. Chi-squared statistic: 28.106. P-value: 1.15*10^−07^ **(d)** Number of studies sharing code and data across all studies. Chi-squared statistic: 44.16. P-value: 3.02*10^−11^ **(e)** Code and data sharing status across primary studies. **(f)** Cumulative plot for code sharing status for 453 manuscripts between 2016 and 2021.

The presence of code availability statements was significantly associated with the availability of shared code (Pearson’s *χ*2= 42.79, p =2.72*10^−9^). Of the 453 studies analyzed, 49.89% shared code, with 79.20% of them having a code availability statement alongside the shared code, while 20.79% of articles shared code but lacked a code availability statement. In contrast, 43.49% of manuscripts shared neither code availability statements nor code, and 6.62% of manuscripts had a code availability statement but did not share any code (Figure 1b). The majority of the manuscripts that provided code availability statements included them in the main text of the manuscript along with the data availability statement, or in the results, or the supplementary section.

### Studies performing secondary analysis were more likely to share the code compared to primary analysis studies

We then separated the articles into studies that generated their own data or primary analyses and studies that relied on already available data or secondary analyses. Our analysis revealed that out of the 204 biomedical research articles that conducted primary analyses, 69.61% shared the raw data (Supplementary Figure 1). Moreover, we found that 36% of the 204 primary analyses had code and data availability statements, while 44.61% had only given a data availability statement and 0.98% only a code availability statement (Supplementary Figure 2). We then investigated the relationship between code availability and the analysis type (primary and secondary analyses) among the 453 articles. We found that 61.66% of the secondary analyses articles shared their code while only 36.27% of primary analyses articles shared their code (Pearson’s *χ*2= 28.106, p=1.15*10^−07^) (Figure 1c).

### Data sharing is a more prevalent practice than code sharing in biomedical research

In this study, we investigated the utilization of data repositories and code repositories to assess the prevalence of code and data sharing practices. Of the 44.64% of biomedical research articles that conducted primary analyses between 2016 and 2021, we found that a significant majority had shared data while not sharing their code (Pearson’s *χ*2= 44.16, p=3.02*10^−11^) (Figure 1d). Specifically, only 29.90% of the articles were accompanied by both code and data. In contrast, 24.02% of articles shared neither the code nor the data. Of the 204 primary analysis manuscripts, 39.71% were accompanied by data only, whereas 6.4% shared code only (Figure 1e).

When we examined the 453 articles on a year-by-year basis, we found that code sharing practices have exhibited a consistent positive trend from 2016 to 2021 with an increase from 7.28% in 2016 to 49.89% (Figure 1f). Data availability appears to be more common than code availability in the field of biomedical research, specifically when looking at primary analysis studies (Supplementary Figure 3). When we looked at primary analyses data year by year, we found that both code and data sharing practices have exhibited a consistent positive trend from 2016 to 2021. Notably, data sharing practices experienced a substantial increase from 9.31% in 2016 to 69.61% in 2021 (Supplementary Figure 4), while code sharing practices also demonstrated a noticeable but significantly slower upward trajectory from 1.99% in 2016 to 16.34% in 2021 (Supplementary Figure 5). In terms of secondary analysis studies, we found that there is a similar upward trend with an increase from 5.30% in 2016 to 33.55% in 2021 (Supplementary Figure 6). In addition to examining the prevalence of data and code availability practices, we also made specific observations regarding the utilization of data repositories.

Our analysis revealed that GEO emerged as the most commonly used repository for sharing data used in analyses, comprising 28.77% of the shared data, followed by SRA at 17.81%. The remaining 47.26% of shared data originated from other repositories (Supplementary Figure 7). Additionally, 65.20% of the articles that shared raw data had a data availability statement mentioned in the manuscript (Supplementary Figure 8). To explore the relationship between the presence of data availability statements and data sharing, we conducted an analysis and found a statistically significant association between the two (Pearson’s *χ*2= 125.13, p=6.07*10-27). This suggests that including a data availability statement in a manuscript can improve access to shared data.

### Positive impact of journals policies on code sharing

In our study, we sought to investigate the differences in code availability policies across eight biomedical journals and examine the impact of these policies on code sharing. We found that out of the eight journals, five mandated code sharing, while three either encouraged code sharing or mandated it upon request (Supplementary Table 1). In contrast, six of the eight journals had a mandatory data sharing policy, and two only encouraged or mandated data sharing upon request. Furthermore, across all journals, we see that 92.27% are sharing their code in non-notebook formats, R Markdown Document (RMD), or Notebook format (Supplementary Figure 9). To investigate the effect of journal policies on code and data sharing, we analyzed the relationship between the journal policy for code and data sharing and whether the actual code and data was available for that article for that specific journal.

For all 453 articles, we found that articles were 0.43 times more likely to share their code if the journal policy is mandatory compared to when it’s encouraged. In contrast, articles were 2.34 times more likely to not share their code if the journal policy only encouraged code sharing (p=1.88*10^−05^). Furthermore, the number of articles whose journals mandated code and shared their code was significantly larger than the number of articles whose journals only encouraged sharing code and shared their code (Pearson’s *χ*2= 17.88, p=2.36*10^−05^). Moreover, when looking specifically at journals that only encouraged code sharing, we found that the percentage of code unavailable was much higher than the percentage of code available (Supplementary Figure 10).

However, when we narrowed our analysis to only primary analyses articles, we found that articles were 0.34 times more likely to share their code if the journal code sharing policy was mandatory compared to when it was only encouraged. On the other hand, articles were 2.93 times more likely to not share their code if the journal policy only encouraged code sharing (p=1.80*10^−3^). Additionally, there was a higher percentage of code not shared across both types of journal policies (Pearson’s *χ*2= 9.24, p=2.37*10^−3^). Nevertheless, the percentage of code available was much lower than the percentage of code unavailable for journals that only encouraged code sharing (Supplementary Figure 11).

When looking at the effect of journal policies on data sharing, we found no significant effect despite the fact that journals that had mandatory data sharing policies had a much larger percentage of code shared than journals that only encouraged code sharing (Pearson’s *χ*2= 1.05, p=0.31). However, in both categories, journals that had mandatory policies and journals that encouraged data sharing, the percentage of articles that shared data was higher than articles that did not share data (Supplementary Figure 12).

The results of our analysis suggest that journal policies can have a significant effect on the availability of code. Journals that require code are more likely to publish papers with code available. This is important because code sharing can help to improve the reproducibility of scientific research.

### Feasible actions to increase availability of analytical code

When journals implemented policies to ensure code and data availability unless there are ethical or legal restrictions preventing it, an increasing number of researchers followed the rules^15^. An analysis of studies utilizing Natural Language Processing revealed that a journal’s introduction of a new policy requiring code sharing resulted in a rate of 53% in 2019, 61% in 2020, and 73% in 2021. Following the introduction of this policy, the code-sharing rate increased to 87% in 2022^6^. While many journals have implemented policies to ensure code and data availability, the current efforts primarily rely on individual researchers. However, there are two main ambiguities in the policies of some journals. First, it is unclear whether “supporting data” and “underlying data used for analysis” refer only to raw data or also include software source code, algorithms, and code used for analysis. Second, it is unclear whether the policy includes making all code used for analytics available or only software source code or custom code. Efforts have been made by the biomedical community to establish effective strategies and best practices to enhance reproducibility and rigor in research such as Code Ocean, a cloud based reproducibility platform that has been implemented by a number of journals including, *GigaScience* since 2017^16^, and since 2020, several Nature journals, BMC Bioinformatics, Scientific Data, and Genome Biology^17^. Sharing code and ensuring reproducibility has extended beyond the non-biomedical publishing space; the *Political Analysis* journal wraps every papers’ methods in Code Ocean as standard^18^. Since 2012, the pioneering open science journal, *GigaScience*, has been mandating all papers to have open data, supporting metadata and source code with Open Source Initiative (OSI) - approved licenses^19^. The journal also has its own repository, GigaDB^20^, where authors can host data (not already open in a community approved repository, e.g., NCBI), in the public domain under a CC0 license. To ensure reproducibility and stability of source code, the GigaDB Curators also take snapshots of all code and a dataset is produced with a citable Datacite DOI (Digital Object Identifier). There are new publishing efforts that aim to address transparency and reproducibility in research by moving beyond static publications. One new journal, *GigaByte*^21^ publishes papers with embeddable and interactive features, such as 3D-image viewer via SketchFab, interactive maps, videos, Hi-C data figures, and methods via Protocols.io. In addition to publishing the epitopepredict^22^, GigaByte also published a Stenci.la^23^ Executable Research Article that enables readers to see and play with the code underlying the supporting figures. The epitopepredict tool was also wrapped in a Code Ocean capsule and embedded in the online article, allowing readers to interact and test the code, as well as deploy it to their own AWS Cloud compute service. *eLife* has also leveraged the Stenci.la platform as part of their Reproducible Document Stack initiative^24^.

This is all part of a larger effort to improve the reproducibility of scientific research. CODECHECK is a great scheme that tackles one of the main challenges of reproducibility by providing guidance, workflows and tools that enables anyone (Codecheckers) to independently test the execution of code underlying research articles^25^. Some journals have already adopted this scheme, where a citable, time-stamped CODECHECK certificate is issued, and credit also given to the Codecheckers^26^. CODECHECK is an easy scheme any publisher and journal can use to ensure reproducibility of papers, and is also a great learning tool for students or anyone interested in data science. Nevertheless, an ongoing dialogue between journal publishers and researchers is necessary to overcome perceived and technical barriers to transparent and reproducible data-driven research across various biomedical disciplines.

Data-driven biomedical research relies heavily on computational analyses, yet there is currently a lack of effective mechanisms to ensure transparency and reproducibility of data analyses performed to generate results in scientific publications. Despite efforts to increase transparency and reproducibility, many published studies remain non-reproducible due to the lack of code and data availability^27^. One reason for this is the researchers’ concerns about losing priority over their novel results, misrepresentation of their work, maintaining exclusivity of their data for future projects, or a fear of data parasitism^28^, as well as lack of credit for making data and source code open. These concerns can be addressed by including the primary authors as co-authors on any publications resulting from previously shared code and data^29^, as well as citing software correctly by citing the article describing the software and the source code^30^.

To address all of these challenges, code and data sharing should be considered an integral part of study submission, similar to ethical review and publication of study results. They should also be considered as early as possible when developing research projects, and sharing policies for studies should be conceived as part of the study design. The neuroimaging community are strong adopters of open science and have suggested four important aspects of scientific research practice: respect trademarks, clarify ownership by looking at copyright status, release code and data under free and open licenses, and finally obtain the permission to share the data or code^31^. In addition, Stanford Data Science have released their Stanford Open By Design Handbook, targetted to early career researchers who want to adopt open science practice, but lack the knowledge and expertise^32^.

Journals and funding bodies should require authors to make raw data available by publishing it on public servers and committing to long-term availability of their code and data. Universities should also provide open science guidelines and support for their researchers. By implementing such practices, researchers can enhance transparency, reproducibility, and the overall quality of data-driven research in the biomedical field.

## Discussion

Despite the growing recognition of transparency and reproducibility in biomedical research, there is no consensus on actionable steps for the scientific community when it comes to data and code sharing, elements that are essential for the advancement of science^33^. Our study provides a comprehensive analysis of code availability in biomedical research, highlighting the importance of sharing code and data. We propose that sharing code and data should be integral to study submission, similar to ethical review and publication of results. Journals and funding bodies should mandate code and data sharing as requirements for publication and funding. By doing so, they can inform and encourage the biomedical community to adopt effective strategies for improving transparency and reproducibility in data-driven biomedical research. To address the obstacles hindering data and code availability, we aim to raise awareness and promote open dialogue among journals, funding agencies, and researchers.

One key benefit of making code freely available is its time-saving aspect, enabling researchers to adapt existing code and avoid starting from scratch^11^. Code sharing fosters collaboration within and across research groups^34,35^, leading to increased efficiency and novel discoveries. Moreover, code availability increases trust and enables researchers to reproduce and validate each other’s results, enhancing the credibility of scientific research. Reusing code across various projects contributes to the development of standardized analytical tools and workflows, further improving research practices^36^.

While data sharing became widely enforced, guidance on code sharing remains limited^37,38^. This imbalance highlights the need to develop comprehensive guidelines and standards that encourage researchers to share code along with their research results^38^. Practices promoting the open sharing of code products and protocols significantly contribute to reproducibility, facilitate secondary analysis, and prevent unnecessary barriers in republishing previous findings^38^.

To foster this collaborative and efficient research environment, the scientific community must promote open science practices, establish clear standards for code sharing and citation, and develop guidelines for reporting preclinical research data^37,38,39,40,41^. However, studies indicate that supporting data reporting in research publications remains inadequate^37,42^. For instance, only 38.1% of immunogenomics research publications shared their data on public repositories^43^. Therefore, it is important to develop and adopt specific guidelines for sharing code products used in analysis and visualizations, similar to those for data sharing^38^. Even so, the debate on how scientists should go about sharing code remains unresolved.

Typically, researchers will share their code using open-source repositories like GitHub. However, relying on these platforms poses challenges, including limited file capacity and concerns about archival stability due to unstable URLs over time. Despite these challenges, the advantages of open-source repositories and URLs currently outweigh the disadvantages, making them a practical option. Nonetheless, the scientific community must engage in a dialogue about the long-term implications and explore alternative solutions to ensure the accessibility and stability of shared code and data in biomedical research.

Sharing research data and code is fundamental for advancing scientific knowledge, enabling collaboration, and accelerating progress. It prevents duplication, allows building upon existing work, and generates critical new findings in the biomedical field and other domains. Openly sharing research data and code ensures the reusability of studies, promotes transparency, and enhances the credibility of research findings^37,38,43,44^. Several recommendations and agreements are already in place for sharing data, such as the Fort Lauderdale agreement in 2003 that reaffirmed the 1996 Bermuda Principles^45^, the Toronto agreement in 2009^46^, and data sharing recommendations by the National Academies^47^. Complementing these efforts, there exists hands-on education initiatives, such as Software Carpentry, which teaches fundamental lab skills for research computing^47,48^. However, there is no single agency responsible for enforcing compliance with these guidelines and procedures. This is a critical oversight, as it means that there is no one entity ensuring the usability and reproducibility of shared data and code. Therefore, it is crucial to identify a responsible agency to fill this role. The community should work towards establishing actionable strategies that align with the Findable, Accessible, Interoperable and Reusable (FAIR) guiding principles^41,49^ to accelerate the sharing of research products. By merging efforts to enhance transparency and reproducibility in biomedical research, we can overcome these challenges and pave the way for more robust and reliable scientific findings.

### Box 1.

Actions to increase availability and archival stability of analytical code

Here, we present five principles to increase code availability and archival stability.

1. **Code should be available and well organized** The most important step to take towards ease of availability is to be organized. By doing some simple steps, we can organize code and data of the project, such as: Data and code should be stored separately. Writing ReadMe files with detailed instructions on how code should be run, dependencies, environment setup, etc. Choosing file names carefully.
2. **Software and resources should be hosted on on archivally stable services** Selecting the appropriate service to host your software and resources is critical. A simple solution is to use web services designed to host source code (e.g., GitHub or SourceForge). To facilitate simple installation, provide an easy-to-use interface to download and install all required dependencies. Ideally, all necessary installation instructions should be included in a single script, especially when the number of installation commands is large. Package managers such as pip or anaconda can potentially make this problem easier to solve.
3. **Journals should incentivize authors to make code publicly available**. Code and data sharing should be considered the main part of submitting a study, similar to how we consider ethical review and publication of study results. Journals and funding bodies should insist that authors make raw data and other supporting data available. Moreover, they should follow recommendations of the National Academies regarding data sharing and abide by the guidelines of the Fort Lauderdale and Toronto agreements^47^. Journals should mandate clear source code and data availability sections to be included in all manuscripts, where accessions for raw data in community-approved repositories are made clear, and other data supporting is cited. Other supporting data should be made openly available in a public server (eg. Zenodo or Figshare) and authors must enter a commitment that their code and data will remain for a long time in stable software archives (e.g., Github, Sourceforge, Bioinformatics.org). Another proposition is implementing a badge system as a simple and cost-effective method to encourage researchers to adopt open practices, thereby increasing transparency in research and promoting the principles of the open science movement ^50^. Nevertheless, a clear policy for sharing code underlying research findings is needed.
4. **Training, seminars and conferences should be held on code and data sharing** Holding seminars and conferences about the advantages of sharing data and code, including hands-on workshops, such as Software Carpentry. PLOS Computational Biology journal presented a new code-sharing policy in March 2021. This policy focuses on improving code sharing. This policy increased repeatability of studies after 2021.

## Methods

### Download and prepare scientific publications for analysis

We downloaded the full text of 12,603 publications between the years 2016 and 2021 from the PubMed Central (PMC) open access corpus in Extensible Markup Language (XML) format from 8 journals: Nature Biotechnology, Genome Medicine, Nature Methods, Genome Biology, Bioinformatics, BMC Bioinformatics, Nucleic Acid Research, and Nature Genetics. We included the commercial use and non-commercial use subsets. From these 12,603 manuscripts, 10 manuscripts were randomly selected from each journal for each year between 2016 - 2021, totaling to 480 randomly selected articles. Each publication had a unique PMC identification number (PMC ID). The XML file for each publication was stored in the corresponding directory based on the journal the publication came from. To further evaluate these manuscripts for sharing the code and data, we filtered out 27 manuscripts which were published as research letters, reports or editorial etc. In total, we had 453 manuscripts available for downstream analysis.

### Extract publish dates of selected publications

We created a key file with the paths to every XML file representing a publication. We then used a Python script utilizing the package *ElementTree* to systematically extract publication metadata from these XML files. To find the date the publication was published, our script considered each publication date that was listed in a publication’s file (there were often multiple). When the year was missing from a date, we disregarded the date completely. When the month was missing, we recorded the publication month as December. When the day was missing, we recorded the publication day as the last day of the publication month. After completing this process with each listed date, the earliest converted date was recorded and associated with our record of the publication.

### Analysis of extracted information

We also recorded various parameters such as the raw data availability, where the data is shared, the analytical code availability, code availability in supplementary files, code availability in repositories or webpages, presence of code and data availability statements, and the number of citations for each study. All parameters recorded were based on information provided by the authors in the publication. Amongst the 480 studies, we filtered out the studies that were conducting secondary analyses while looking into raw data availability. When we looked into the raw data availability in each article, we assigned ‘Yes’ and ‘No’ to indicate availability and unavailability of raw data. The data was considered available if the data and its corresponding dependencies could be stored on public repository operating systems, and the data can produce expected results from the input data with no errors. We also recorded whether the data was shared on ‘Gene Expression Omnibus (GEO)’, ‘Sequence Reads Archive (SRA)’ or other repositories such as Bioproject, ENCyclopedia Of DNA Element (ENCODE), DNA Database of Japan (DDBJ), GitHub, ArrayExpress, European Nucleotide Archive (ENA). Those articles which shared data on either DDBJ or ENA were included under ‘SRA’. Some articles had data shared or obtained from multiple datasets for different parts of their study and these articles were included under the category ‘Multiple’. The category ‘Multiple’ not only refers to articles which have split their data on multiple repositories but also could refer to those which have duplicated datasets on different repositories.

Similarly, we recorded the code availability in all the 480 articles and assigned ‘Yes’ or ‘No’ to indicate availability and unavailability of analytical code. The goal of this study was not to check the completeness of analytical code but the presence and incentive of researchers to share code. We specifically looked for code used for analysis (not to be confused with software or source code for algorithms and tools) and considered code availability as a ‘Yes’ if any analytical code was shared. For those articles that shared code, we recorded if the code was hosted on GitHub, shared in the supplementary materials or hosted other repositories such as Zenodo or a webpage. Some articles that shared code in both supplementary materials and on Github were included under GitHub. Verifying the completeness and/or accuracy of the code shared was beyond the scope of this study. Additionally, we also recorded whether the articles had data availability statements and code availability statements mentioned in a separate section for easy accessibility. We assigned a ‘No’ if the article did not have a statement but mentioned the availability of data or code amidst the main text. We also recorded the number of citations the articles had on Google Scholar as of 15 November, 2021, and the policies of the ten journals on sharing code and data. We classified the code and data sharing policies of the journals as ‘Mandatory’, ‘Encouraged/Mandatory’ if it was unclear, and ‘No policy’ for those that did not mention any guidelines for sharing code or data.

### Statistical Analysis

Chi-square tests of independence were used to identify significant differences between groups in most of the analyses. To specifically determine whether journal policies had an effect on code and data availability, the standard odds ratio was calculated and a Fisher’s exact test was conducted to determine whether the odds ratio was significant. All statistical tests were carried out using scipy 1.11.1 package.

## Data Availability

All manuscript data discussed in this study is freely available at https://github.com/Mangul-Lab-USC/code-availability. This data was used to produce all results and figures depicted in this study.

## Code Availability

All code required to produce figures and analysis performed in this study is freely available at https://github.com/Mangul-Lab-USC/code-availability. The code is distributed under the terms of the General Public License version 3.0 (GPLv3). Source data are provided with this study as described in the Data Availability section.

