## Supplementary Figures & Tables for "Analytical code sharing practices in biomedical research"

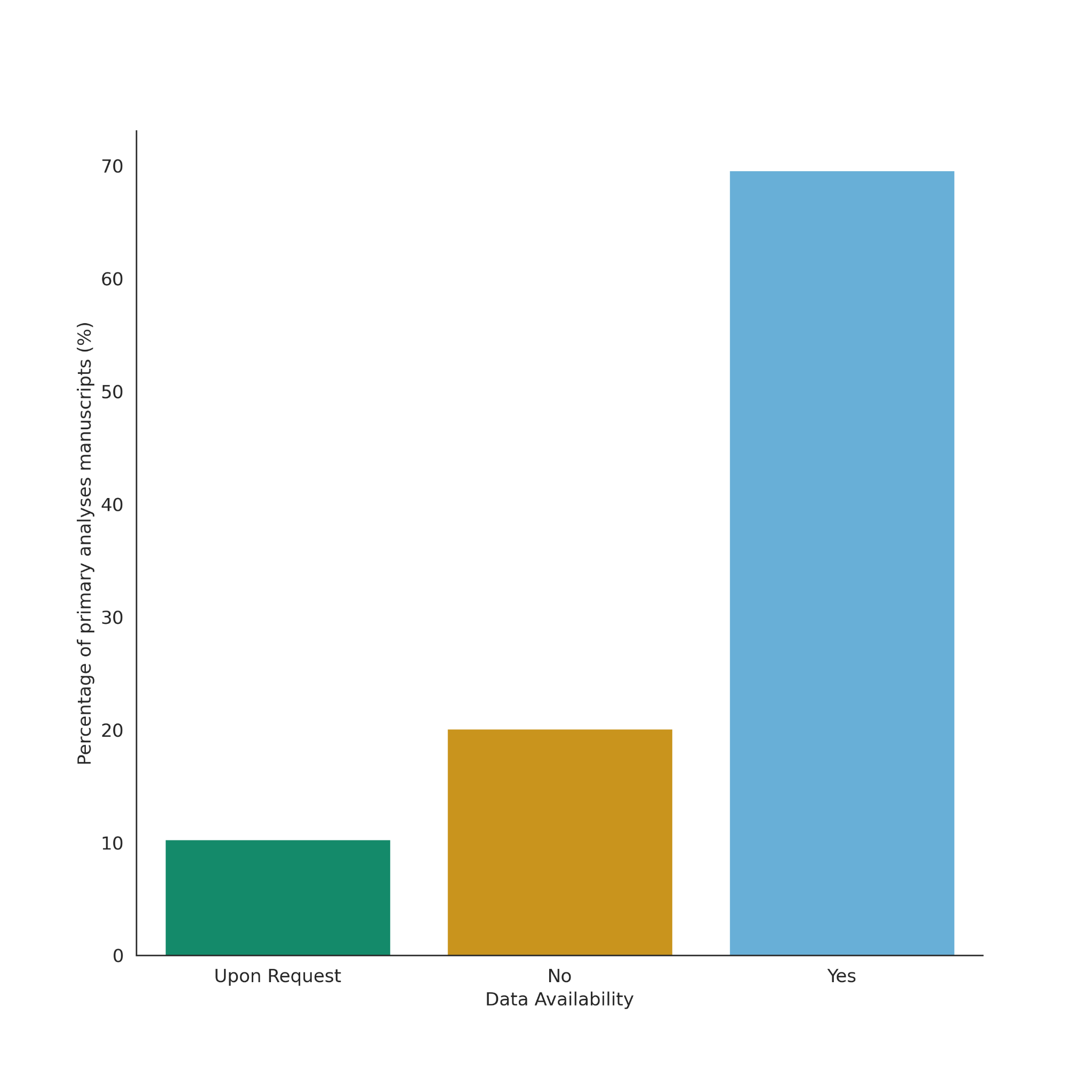

**Supplementary Figure 1**: Data availability status in the primary analysis studies.

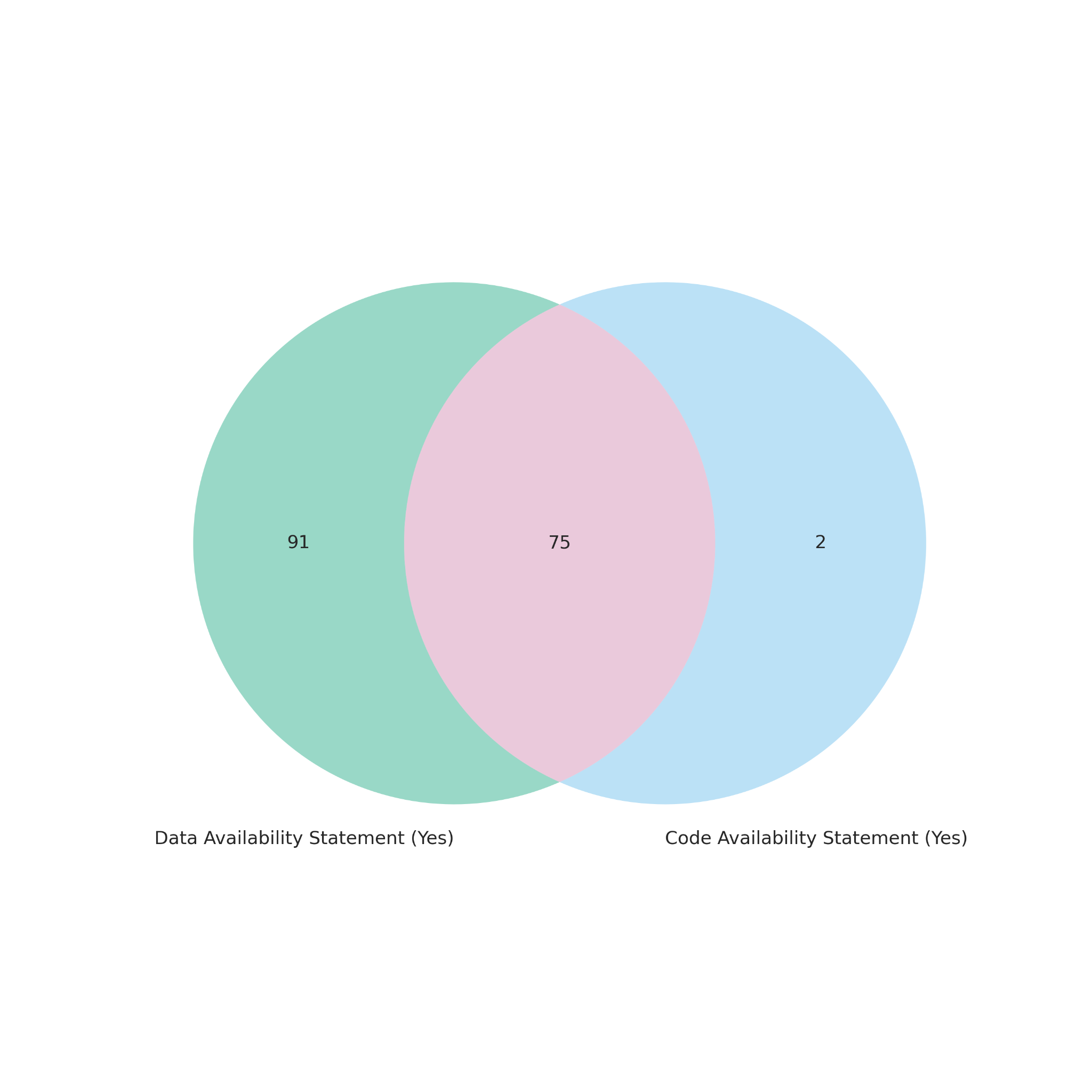

**Supplementary Figure 2**: Manuscripts overlap of code and data availability statement for primary data analysis studies.

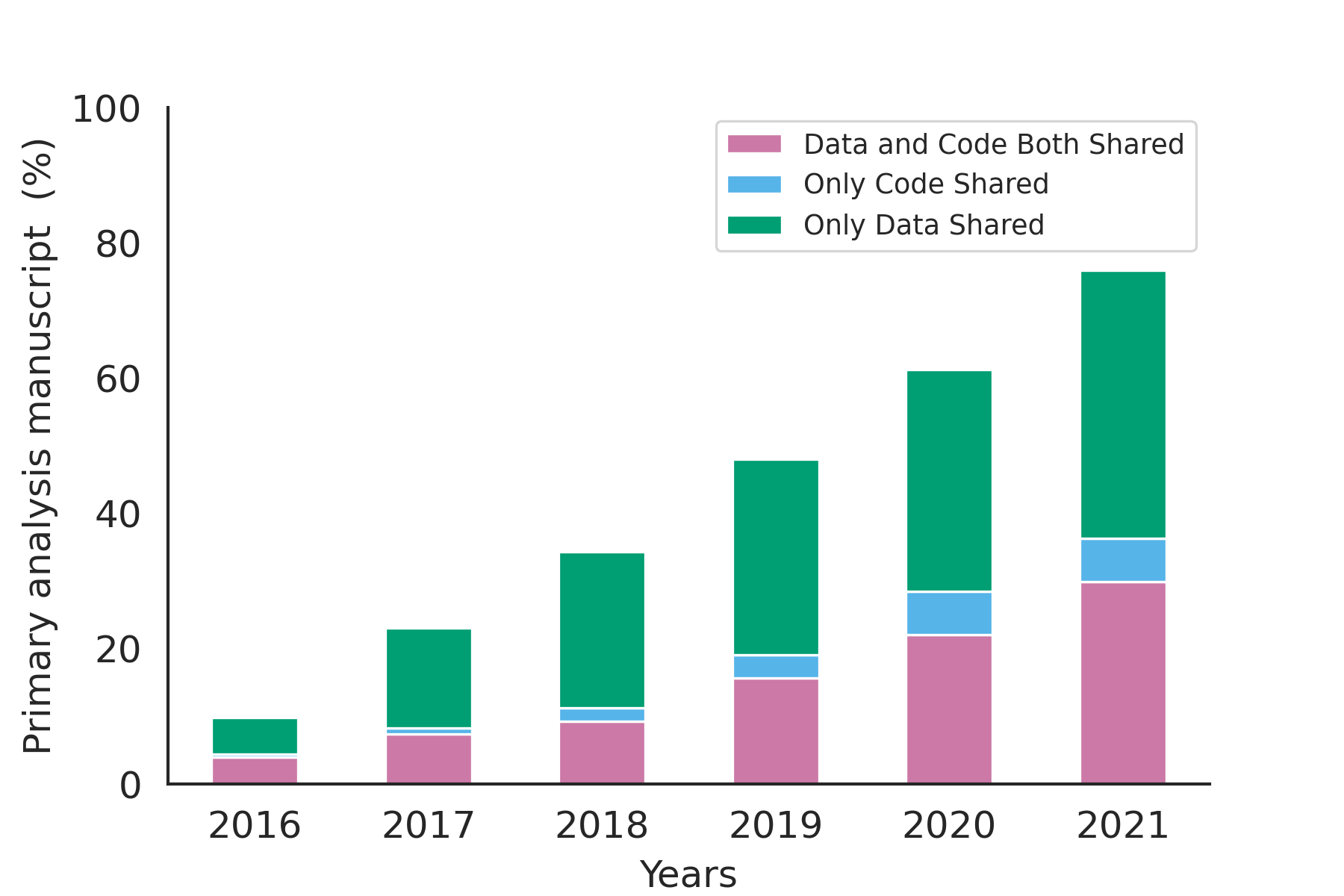

**Supplementary Figure 3**: Cumulative plot for data and code sharing for primary analysis manuscripts between 2016 and 2021.

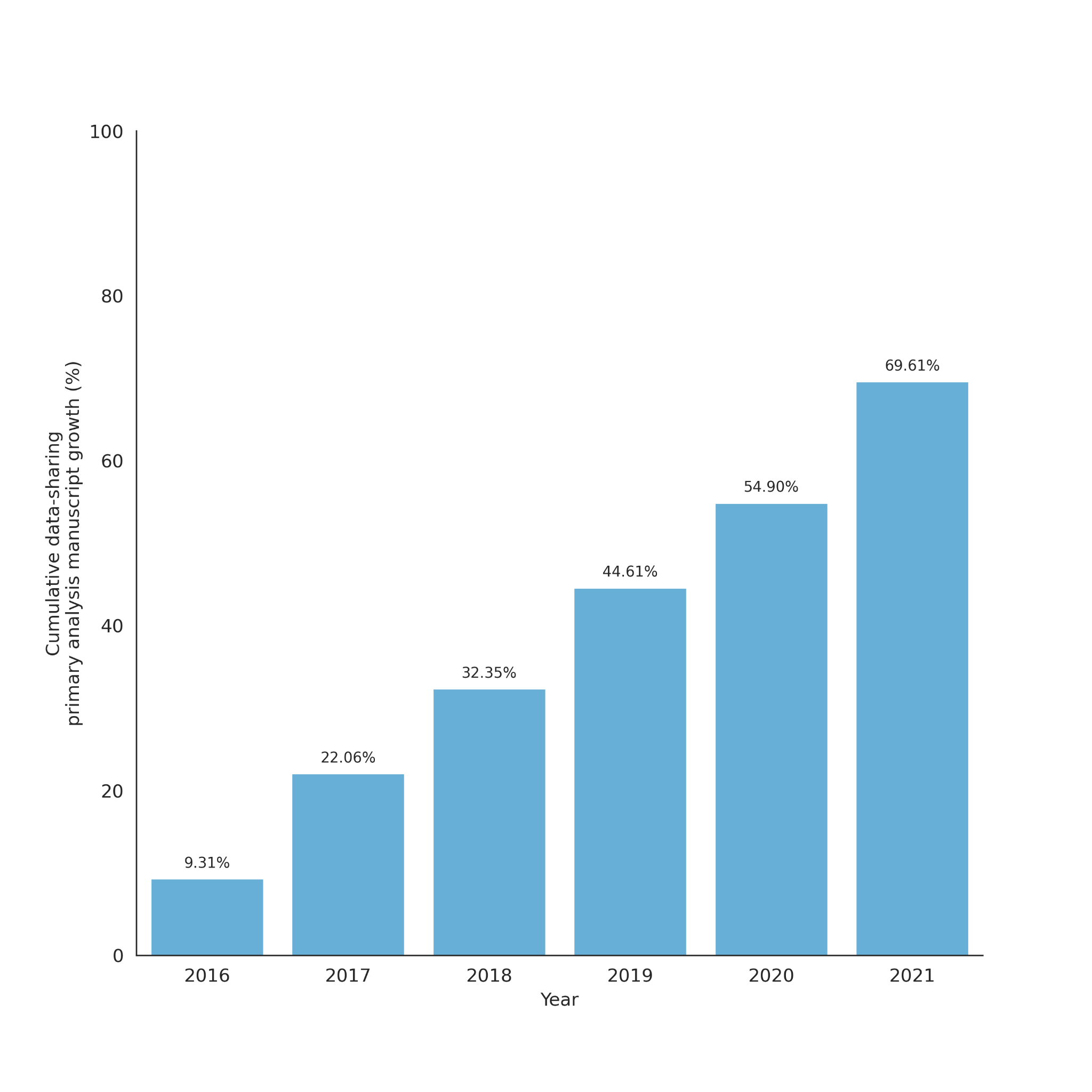

**Supplementary Figure 4**: Cumulative plot for data sharing status for primary analysis manuscripts between 2016 and 2021 (n=204).

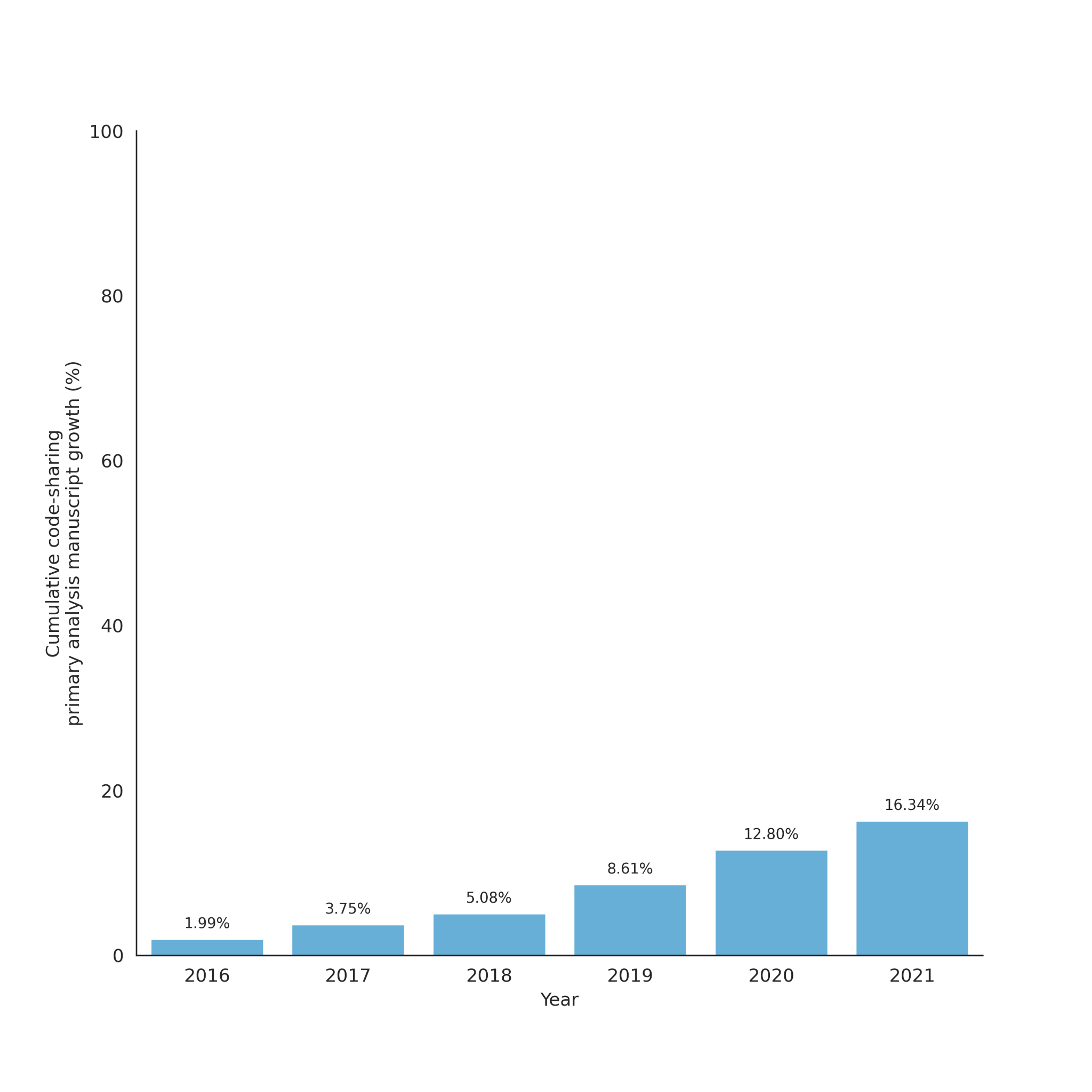

**Supplementary Figure 5**:Cumulative plot for code sharing status for primary analysis manuscripts between 2016 and 2021 (n=204).

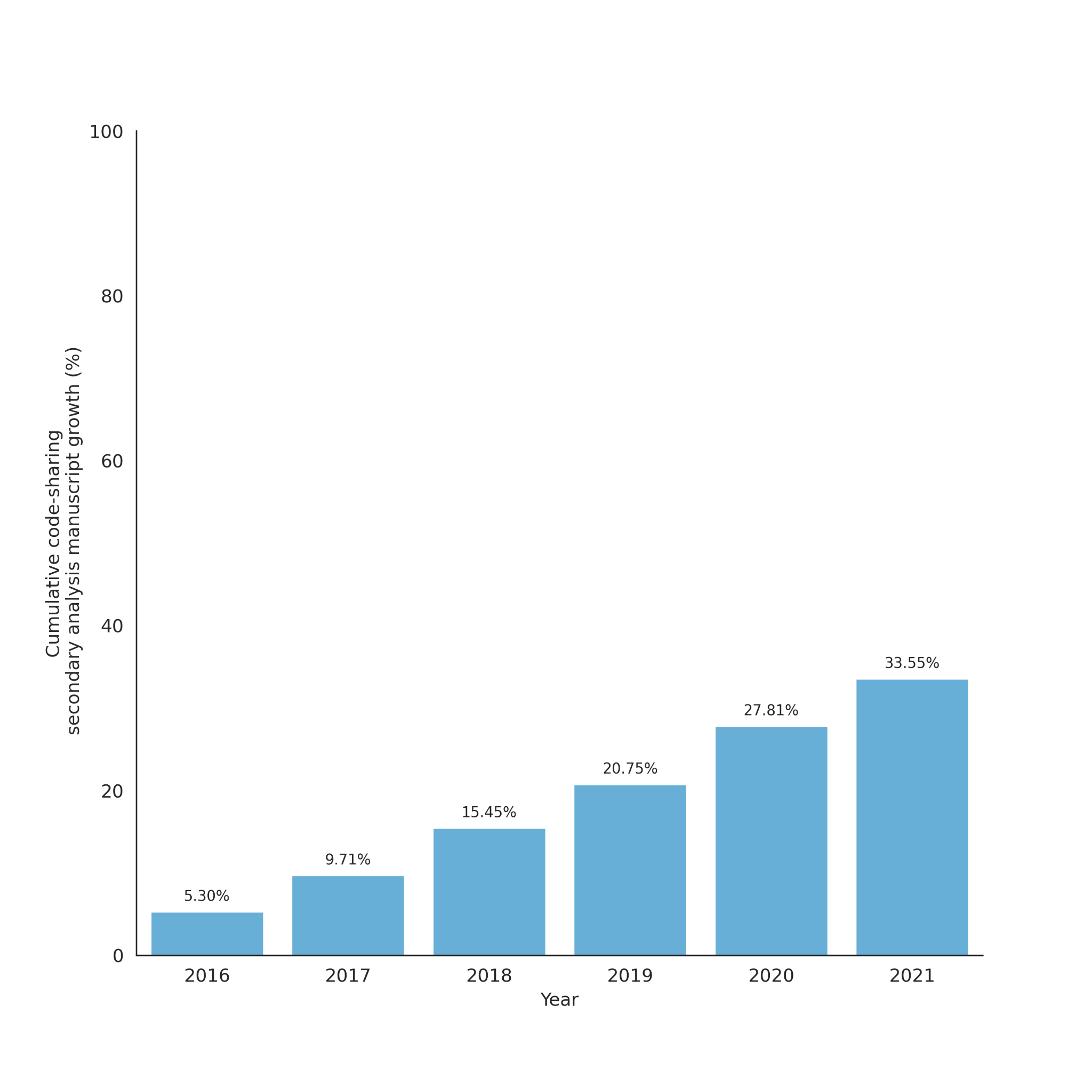

**Supplementary Figure 6**: Cumulative plot for code sharing status for secondary analysis manuscripts between 2016 and 2021 (n=249).

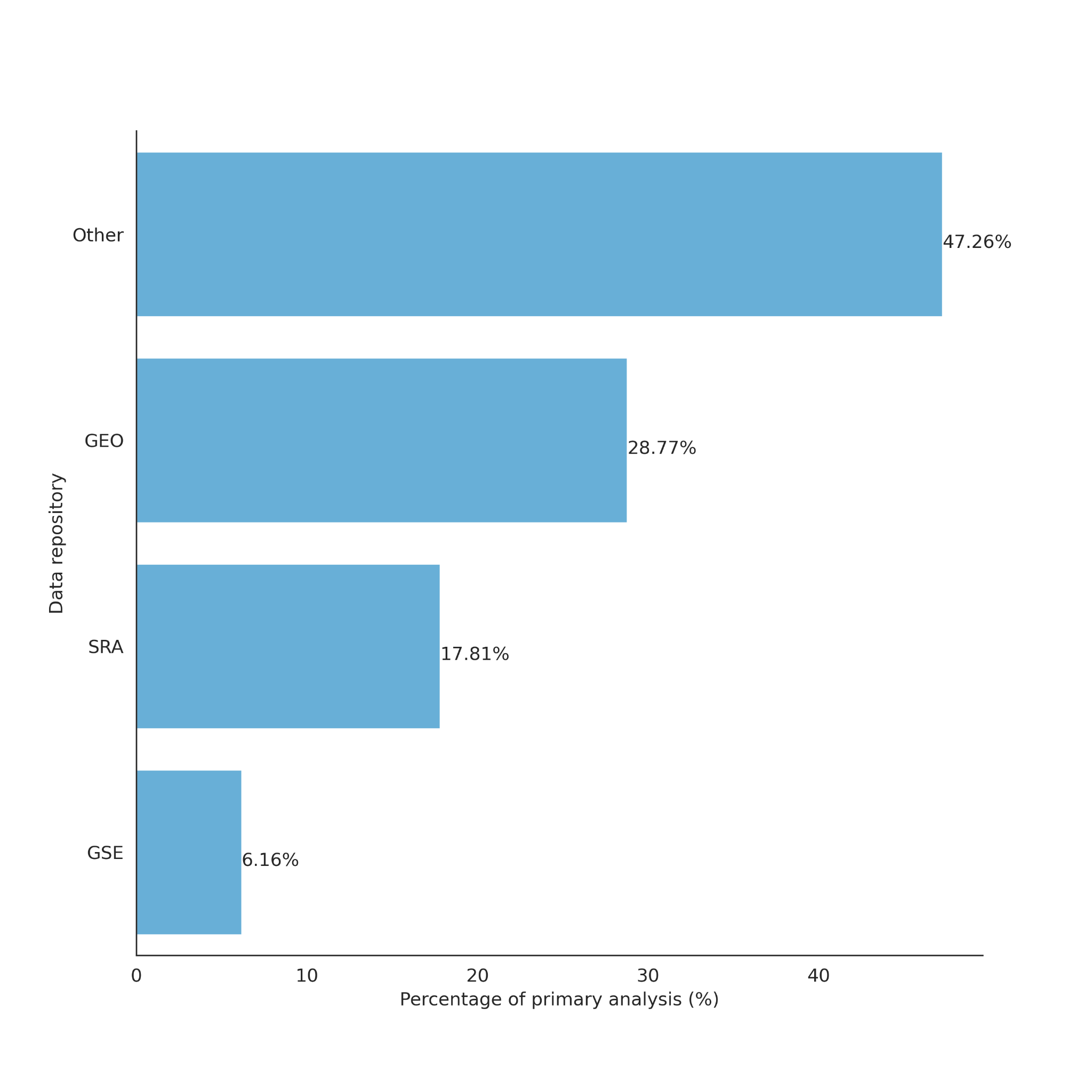

**Supplementary Figure 7**: Data sharing repository for primary analysis studies (GEO:Gene Expression Omnibus, SRA: Sequence Read Archive, GSE: Gene Expression Omnibus).

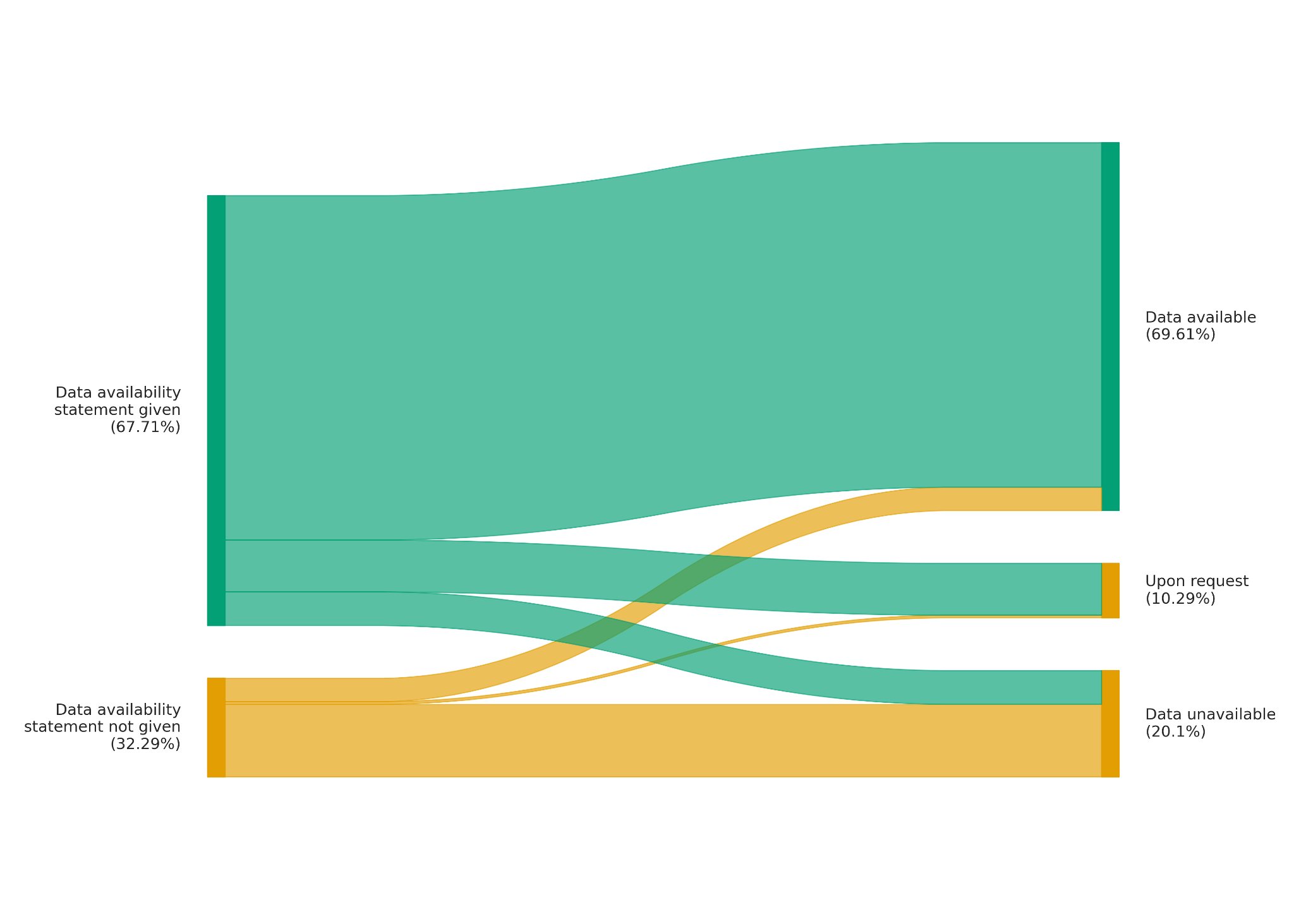

**Supplementary Figure 8**: Data availability status across manuscripts on the basis of data availability statement.

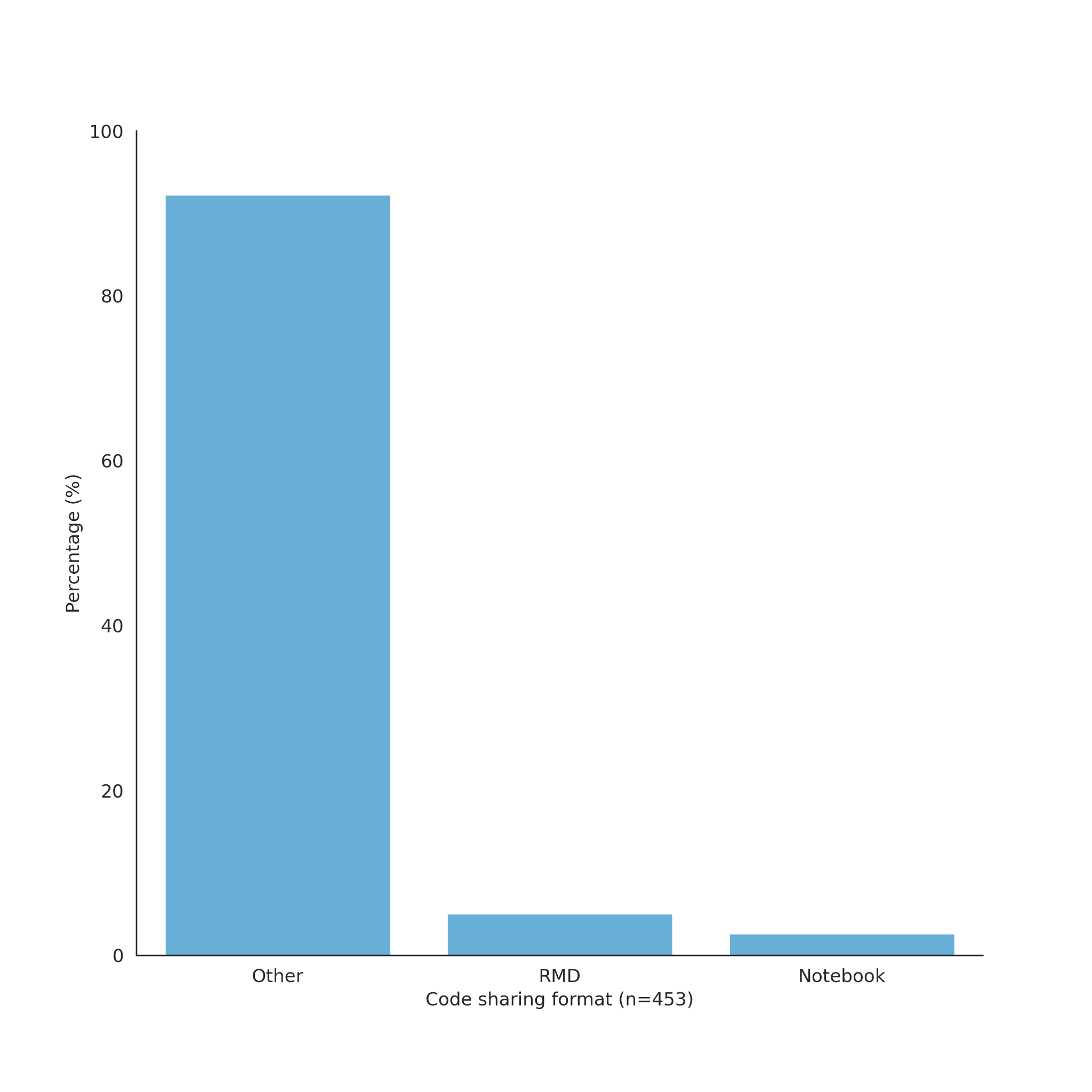

**Supplementary Figure 9**: Percentage of studies that shared code in the following formats: Rmd, notebook or no, which means that no code was shared (n=453).

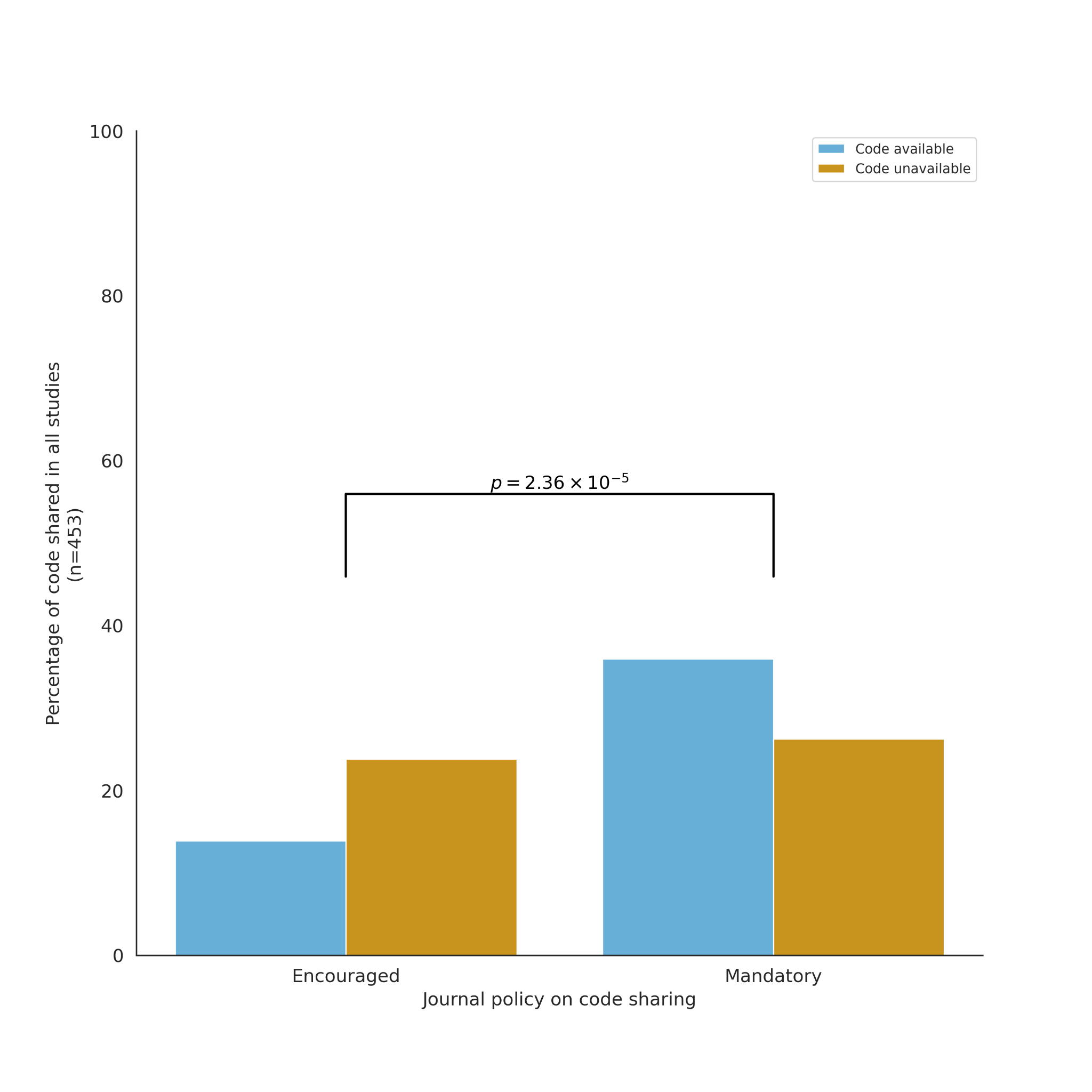
**Supplementary Figure 10**: Code sharing in all studies across 8 journals, by journal policy (n=453).

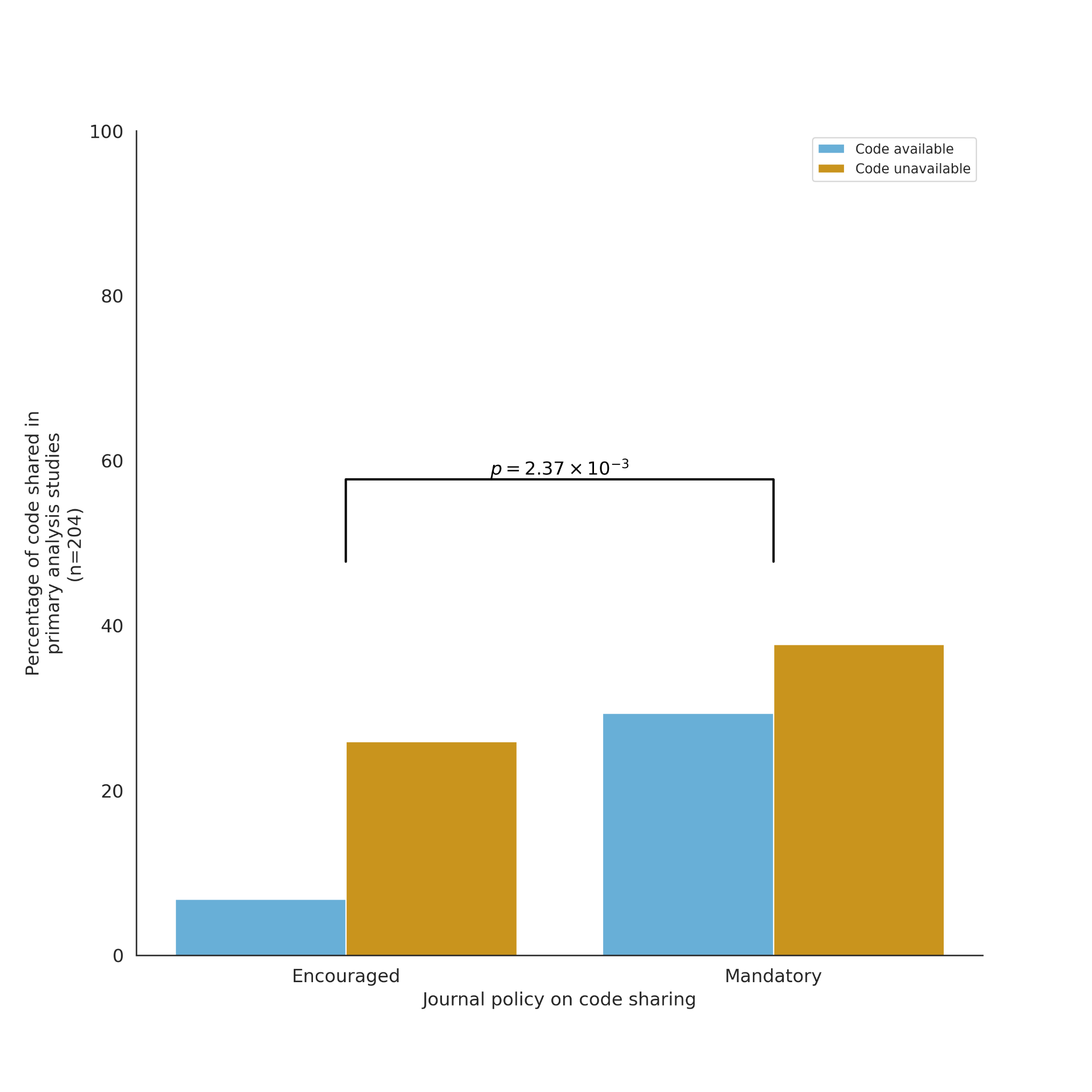
**Supplementary Figure 11**: Code sharing in primary analysis studies across 8 journals, by journal policy (n=204).

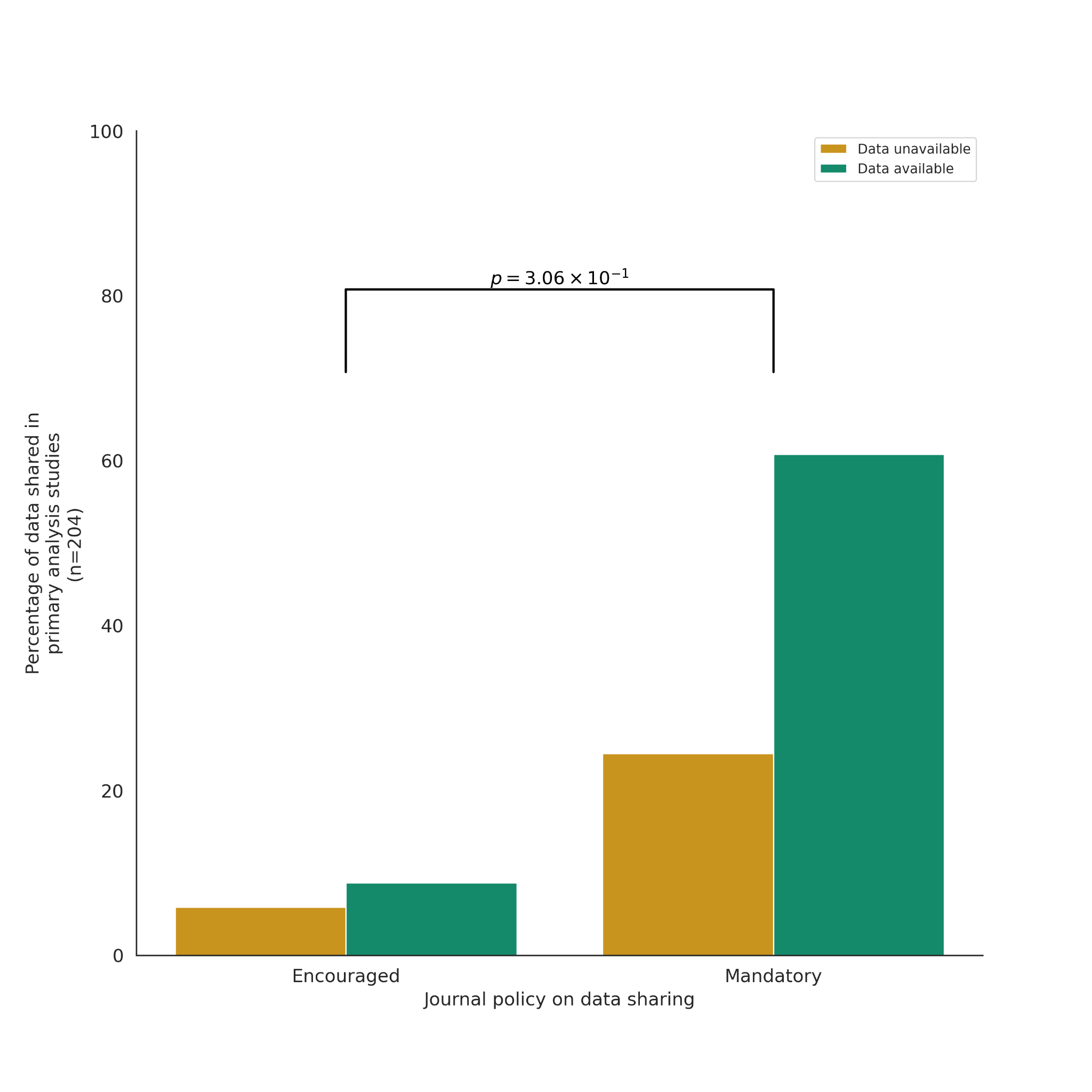

**Supplementary Figure 12**: Data sharing in primary analysis studies across 8 journals, by journal policy n=(204).

**Supplementary Table 1**: Availability of code across 453 biomedical articles classified according to the code repository.

|  | **Source** | **Counts** | **Percentage** |
| --- | --- | --- | --- |
| **0** | 10xGenomics | 1 | 0.43 |
| **1** | Downloadable File | 2 | 0.85 |
| **2** | GitLab | 2 | 0.85 |
| **3** | CRAN | 2 | 0.85 |
| **4** | Bioconductor | 2 | 0.85 |
| **5** | Sourceforge | 3 | 1.28 |
| **6** | Bitbucket | 6 | 2.56 |
| **7** | Supplementary | 6 | 2.56 |
| **8** | Zenodo | 14 | 5.98 |
| **9** | Website | 16 | 6.84 |
| **10** | GitHub | 180 | 76.92 |
